# Comprehensive profiling of TCRβ repertoire in a non-model species (the bank vole) using high-throughput sequencing

**DOI:** 10.1101/217653

**Authors:** Magdalena Migalska, Alvaro Sebastian, Jacek Radwan

## Abstract

In recent years, immune repertoire profiling with high-throughput sequencing (HTS) has advanced our understanding of adaptive immunity. However, fast progress in the field applied mostly to human and mouse research, with only few studies devoted to other model vertebrates. We present the first in-depth characterization of the TCRβ repertoire in a non-model mammal with limited genomic resources available – the bank vole (*Myodes glareolus*). We used 5′RACE and Illumina HTS to describe V and J segments and to qualitatively characterize preferential V–J segment usage and CDR3 length distribution. Finally, a molecular protocol integrating unique molecular identifiers was used for quantitative analysis of CDR3 repertoire with stringent error correction. We found 37 V and 11 J genes that were orthologous to mice genes. A conservative, lower bound estimation of the TCRβ repertoire was 1.7–2.3×10^5^ clonotypes, and the degree of sharing of the observed repertoire between any two individuals was 3.6% of nucleotide sequences and 14.3% of amino acid sequences. Our work adds a crucial element to the immunogenetic resources available for the bank vole, an important species in ecological and evolutionary research. The workflow that we developed can be applied for immune repertoire sequencing of non-model species, including endangered vertebrates.

## Introduction

The central role of T cells in the adaptive immune response^1^, both humoral and cytotoxic, is mediated by T-cell receptors (TCRs), which are responsible for self/non-self distinction and foreign antigen recognition in the context of the major histocompatibility complex (MHC). TCRs are heterodimeric membrane proteins formed by two chains–α and β. The variability of TCRs, necessary to ascertain specificity and robustness for interaction with different antigens, is generated similarly to that of immunoglobulins–that is, by a random, somatic recombination of different, germline DNA segments (Fig. 1). During somatic recombination, V (variable) and J (joining) segments form the variable domain of the α-chain, whereas the equivalent domain of the β-chain is composed of the recombination of three segments: V, J, and D (diversity). In the murine genome, there are approximately 100 V and 60 J segments of α-chain, and 35 V, 12 J, and 2 D segments of the β-chain; however, up to one third of them are pseudogenes or otherwise non-functional^2^. The most variable fragment of a TCR – the Complementarity Determining Region 3 (CDR3) – is responsible for recognition of the peptide-MHC complex. CDR3 encompasses the V(D)J junction, where insertion and/or deletion of random nucleotides between the rearranged segments (N-diversity regions) increases diversity over that introduced by recombination between V, D, and J genes. As a result, each new T cell is provided with a potentially unique receptor, generating an extremely diverse TCR repertoire. But, due to a large random component in CDR3 formation, many TCRs are either non-functional (i.e. not able to bind antigen–MHC complexes) or self-reacting. Consequently, the functional TCR repertoire is shaped during maturation in the thymus, where processes of positive and negative selection assure MHC restriction and central tolerance of T cells, respectively (reviewed in Klein *et al*.)^3^. The size of an individual αβ TCR repertoire of mature lymphocytes has still not been precisely determined even in humans, with estimates differing by an order of magnitude^4–5^ from 2.5 × 10^7^ to 1 × 10^8^. In the house mouse (*Mus musculus*), the number of unique TCR αβ pairings was estimated at 1.9×10^6^, with the number of unique TCRβ chain types estimated to reach 5–8×10^5^clonotypes^6^.

**Figure 1.**
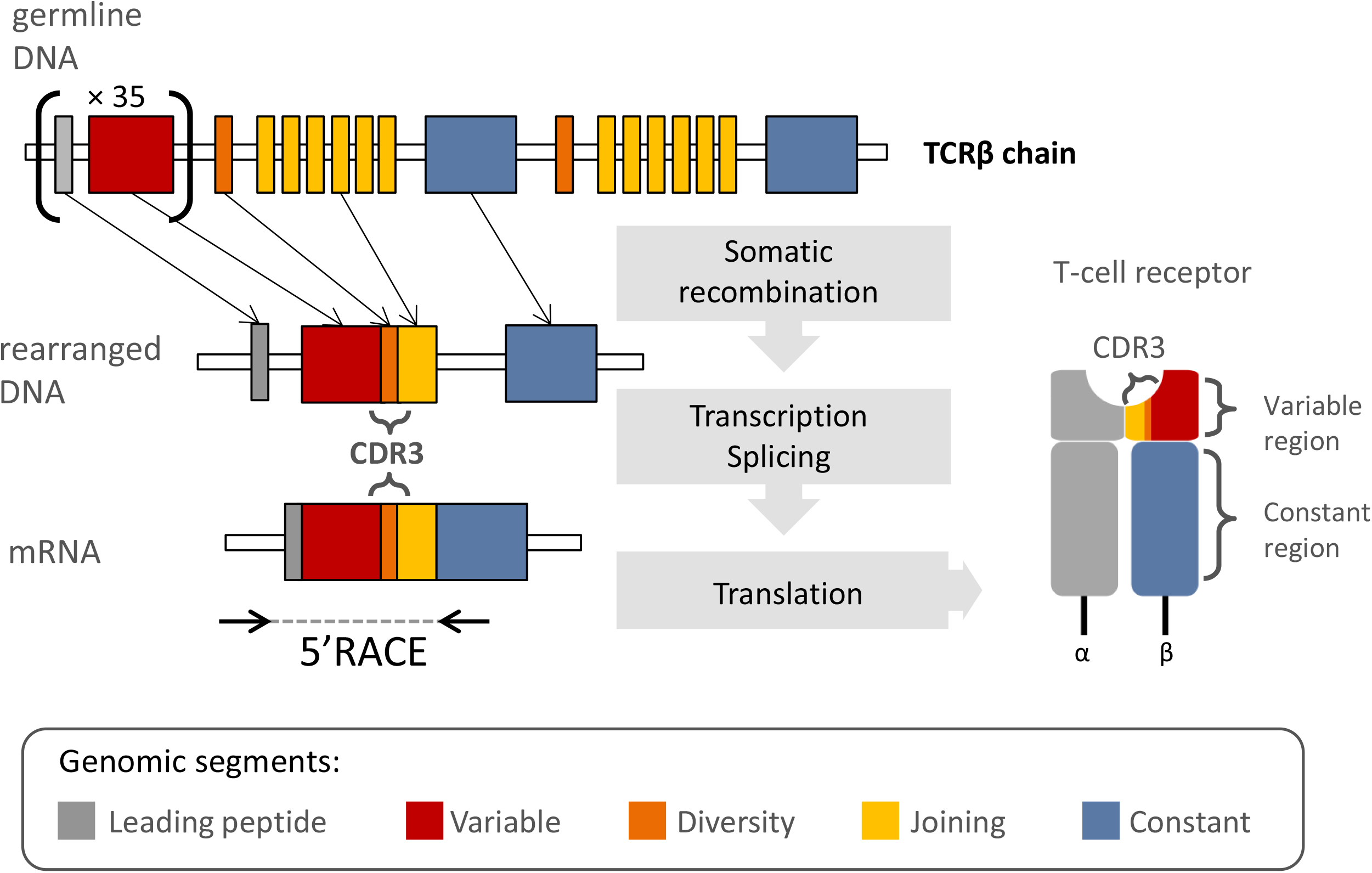
Schematic representation of the TCRβ chain rearrangement. The antigen-binding domain of a TCR is formed by the variable region of each chain, encoded by recombined V, J, and D segments. The Complementarity Determining Region 3 (CDR3) encompasses the VDJ junction with N-diversity regions. Coding TCRβ mRNA is composed of the following segments: L-leading region, V-variable segment, D-diversity segment, J-joining, and C-constant.

The composition and size of the TCR repertoire is crucial for a successful immune response^7–9^; however, its enormous diversity has long impeded in-depth study of an individual TCR’s collections at the genomic level. Indirect techniques, such as CDR3 spectratyping^10^, allowed a general overview of TCR repertoire dynamics, but a direct exploration of sequence diversity was only possible with exceedingly time-consuming and costly cloning and Sanger sequencing (reviewed in Six *et al*.)^11^. Thus, profiling of immune repertoires was limited to a few model species of medical or veterinary importance. However, in the past decade, high-throughput sequencing (HTS) has revolutionized and prompted direct, affordable, and efficient sequencing of immune repertoires^12,13^. The accuracy of TCR repertoire characterization (focusing on the short, but extremely variable, CDR3 region) has further been improved by using 5′ Rapid Amplification of cDNA Ends (5′ RACE)^14^, which alleviates the amplification bias introduced by a multiplexed PCR^15,16^. Further integration of unique molecular identifiers (UMIs)^17^ into the 5′ RACE protocol allows efficient correction of PCR and sequencing errors^18^. The sequence-level resolution of HTS studies has already granted new insights into various aspects of adaptive immunity, such as immune response to infection^19^, autoimmune diseases^20^, repertoire changes with age^5^, or cancer progression and persistence^21,22^. Despite the potential offered by HTS, few attempts were made to transfer these advances to non-model species or, in general, species other than human and mouse. Rare examples of comprehensive analysis of highly variable TCR repertoires with HTS include zebrafish (*Danio rerio*)^23^ and rhesus monkey (*Macaca mulatta*)^24,25^,–both of which are well-established models in medical and biological research, such as embryonic development, viral infections, and vaccine development.

In the present study, we applied recent developments in TCR repertoire sequencing to a non-model mammal, with limited genomic resources available, which is, nonetheless, subject to much ecological and evolutionary research. The bank vole (*Myodes glareolus*) is a small rodent of the Cricetidae family, widespread in western Europe and northern Asia, that is very convenient to study a range of topics–from adaptive radiation^26,27^, genetic basis of adaptation^28^, and response to selection^29^ to processes shaping sexual and behavioural traits^31^. Interest focused on its immunity as well as its role as a reservoir of a pathogenic Puumala hantavirus (PUUV)^32,33^, which causes a mild form of haemorrhagic fever with renal syndrome in humans. To date, a number of components of the bank vole’s immune system have been studied, such as the MHC^34–37^ and Toll-like receptors^38,39^. But, sequencing and profiling of immune repertoires, such as TCRs, has never been attempted in this species.

Here, we adapted protocols available for human and mouse to present the first in-depth characterization of a TCRβ repertoire in a non-model rodent. We took advantage of the fact that the 5′ RACE technique does not require prior knowledge of V and J segments in the species of interest. Necessary primers within a constant region of the TCRβ were designed based on *de novo* assembled transcriptomes–an approach successfully tested before in the highly complex MHC gene family^37^. Amplified transcripts of the TCRβ chains from seven bank voles contained full CDR3 sequences, as well as entire V and J segments. These sequences were subjected to detailed qualitative analysis, which involved description of V and J genes, V–J segment usage, and CDR3 length distribution. For three animals, an estimation of TCR repertoire size was undertaken using a protocol that incorporated UMIs.

## Results

### Qualitative description of the bank vole TCRβ repertoire

Using 5′ RACE and Illumina paired-end 300-bp MiSeq sequencing (Supplementary Fig. S1), we obtained a full variable region of TCRβ chains from seven bank voles, from which V and J segments were extracted. In this section, we describe phylogenetic comparison of these segments with mouse genes, which also served as a basis for naming of the bank vole loci. Furthermore, we analysed V–J segment usage and CDR3 length distribution.

## V and J segment identification and naming

Despite long evolutionary distances between murine and vole sequences, orthologous genomic murine segments could be identified for most J and V bank vole segments (Figs. 2 and 3). Because the bank vole neither has a high-quality annotated reference genome nor are TCR germline gene segments available in the Genbank or International ImMunoGeneTics information system® (IMGT), we assigned J and V segments extracted from TCRβ transcripts to provisional loci based on orthology (Supplementary Fig. S2 and S3) and similarity to the mouse references from the IMGT database^2^ (Supplementary Table S1). Bank vole genes were named BV_TRBV or BV_TRBJ, with the numbering of each locus corresponding to the murine orthologue. With only a few exceptions, there were between one and two variants (hereafter, *alleles*) belonging to each phylogenetic clade (hereafter, *locus*) present per individual (Supplementary Figs. S4 and S5). Such a pattern is expected in a diploid organism and further justifies treating clades as separate loci.

**Figure 2.**
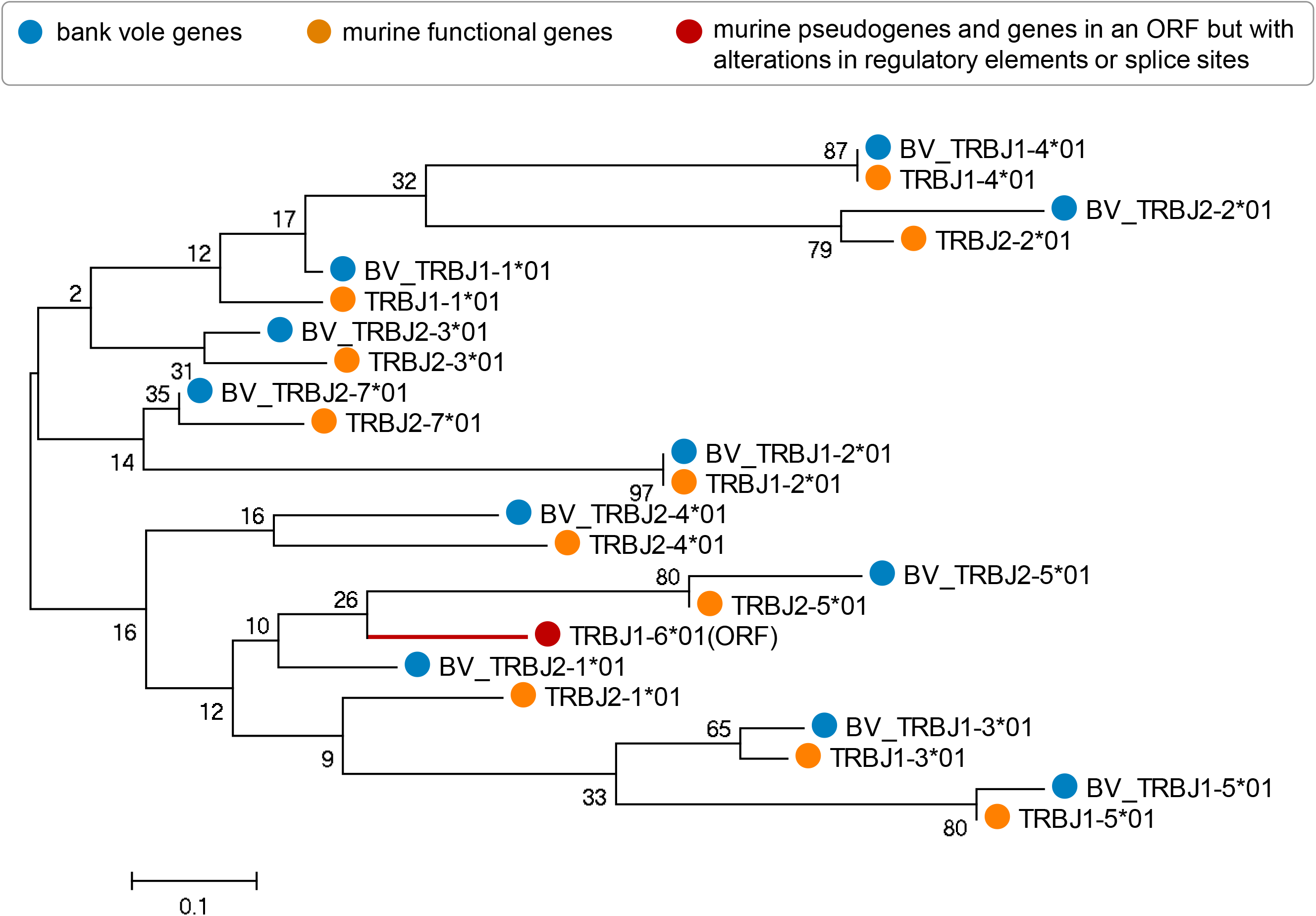
Phylogenetic tree of nucleotide sequences of TCRβ J segments of bank vole and mouse. The tree was inferred using the neighbour-joining method. Bootstrap values based on 1000 replicates are shown. The branch lengths are in units of evolutionary distances (the number of base substitutions per site). The full list of mouse reference sequences with accession numbers is available in Supplementary Table 1. Sequences of bank vole genes identified in this study are available in Supplementary File 1; only one allele for each locus is present herein for clarity. Orange circles mark the mouse sequences (TRBJ*XX*\*01); red circle and tree branch indicates murine sequence in the open reading frame but with alterations in regulatory elements or splice sites; and blue circles mark the bank vole sequences (BV_TRBJ*XX*\*01).

**Figure 3.**
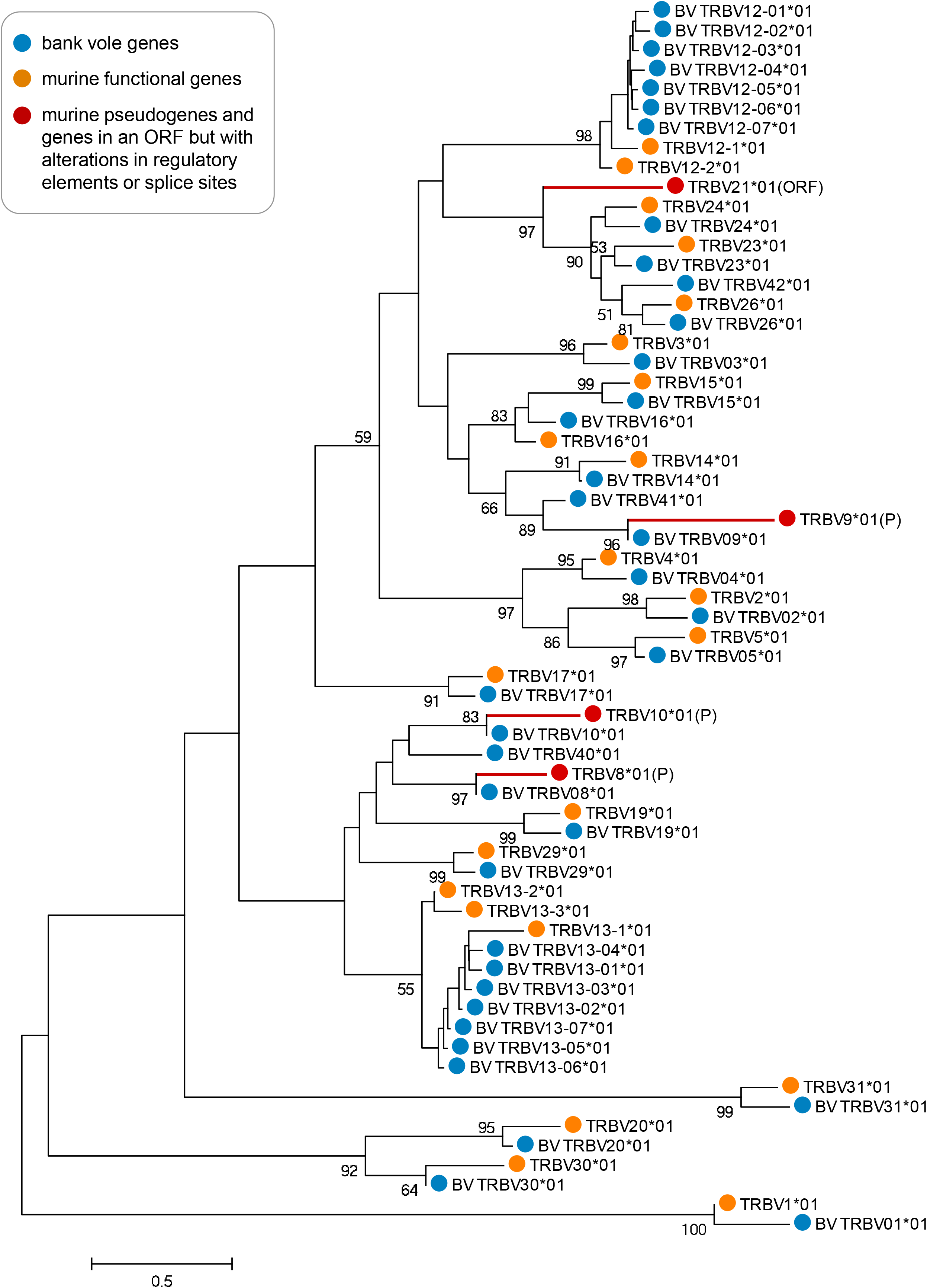
Phylogenetic tree of nucleotide sequences of TCRβ V segments of bank vole and mouse. The tree was inferred using the neighbour-joining method. Bootstrap values greater than 50, based on 1000 replicates, are shown. Branch lengths are in units of evolutionary distances (the number of base substitutions per site). The full list of mouse reference sequences with accession numbers is available in Supplementary Table 1. Sequences of bank vole genes identified in this study are available in Supplementary File 2; only one allele for each locus is present herein for clarity. Orange circles mark the mouse sequences (TRBV*XX*\*01); red circles and tree branches represent murine pseudogenes or sequences in an open reading frame but with alterations in regulatory elements or splice sites; and blue circles mark the bank vole sequences (BV_TRBV*XX*\*01).

We identified 25 distinct V-segment subgroups. Most of them comprised only one locus; however, two subgroups, BV_TRBV12 and BV_TRBV13, contained multiple loci. Similar expansion took place with the murine orthologue subgroups TRBV12 and TRBV13, which have three loci each. Overall, in the bank vole, we found 37 V putative loci (with 1–5 alleles per locus, summing to a total of 116 alleles across loci), compared to 21–22 functional loci in the mouse^40^. In the case of the J segment, we identified 11 loci within two subgroups–one with five loci and the other with six loci. This number exactly matches the number of functional J loci in the mouse^40^. We identified 1–3 alleles per locus, with a total of 17 alleles across loci.

We compared TCRβ V segments retrieved in the present work from directly sequenced transcripts of the bank vole TCRβ against those extracted from a recently assembled draft bank vole genome. These were deposited in the online database Vgenerepertoire.org^41^, containing automatically extracted V genes from whole-genome assemblies (WGA) for over 200 vertebrate species. Only 17 TCRβ V genes were extracted from the bank vole draft genome, compared to 37 putative loci found herein. Comparison of amino acid sequences (Supplementary Fig. S6) showed that the following subgroups of BV_TRBV genes are missing from the WGA: 1, 2, 5, 12, 13, 20, 26, 40, and 42. All of these, except for 40 and 42, have clear murine orthologues, with subgroups 12 and 13 containing multiple duplicated loci. The reason for this discrepancy most likely is the rather poor quality of the bank vole draft genome (scaffolds: 367 242; contigs: 2 834 384; contig N50: 1 591; L50: 365 195), which did not allow successful recovery of all functional genes. All extracted genes from WGA are, however, present in our dataset, with the exception of V49. This locus is likely non-functional and not transcribed, similarly as its murine orthologue TRBV21 (Supplementary Fig. S6), known for alterations in the splicing sites^42^.

## V-J segment usage and CDR3 length distribution

Based on the assignment of the V and J segments of the bank vole to the provisional loci, we analysed preferential usage of V and J segments in the rearranged TCRβ transcripts. In all seven bank voles, the dominant gene segments were BV_TRBV23, BV_TRBV24, BV_TRBV13–1 (Fig. 4a), and BV_TRBJ1–1, BV_TRBJ1–4 (Fig. 4c). Accordingly, combinations of the above segments formed the most favoured V–J pairs, although BV_TRBV23/BV_TRBJ1–4 was the only pairing consistently ranked in the top 1% in all the individuals. Other most common combinations, found in the top 1% of the highest frequency pairs, were BV_TRBV13–1/BV_TRBJ 1–4 and BV_TRBV23/BV_TRBJ 1–1. A comparison of preferred V–J pairs among individuals revealed both between-individual similarities and differences. Figure 5 shows an example of contrasting patterns of V–J gene usage. Individual s02 is characterized by a strong bias toward pairings involving V segment 23 with different J segments from subgroup 1 (BV_TRBJ1–1, BV_TRBJ1–2, and BV_TRBJ1–4) and a very scarce use of J segments from subgroup 2 in all combinations. In contrast, individual s03 has a more even usage of all J segments, including those from subgroup 2. A comparison of all seven individuals is shown in Supplementary Fig. S7.

**Figure 4.**
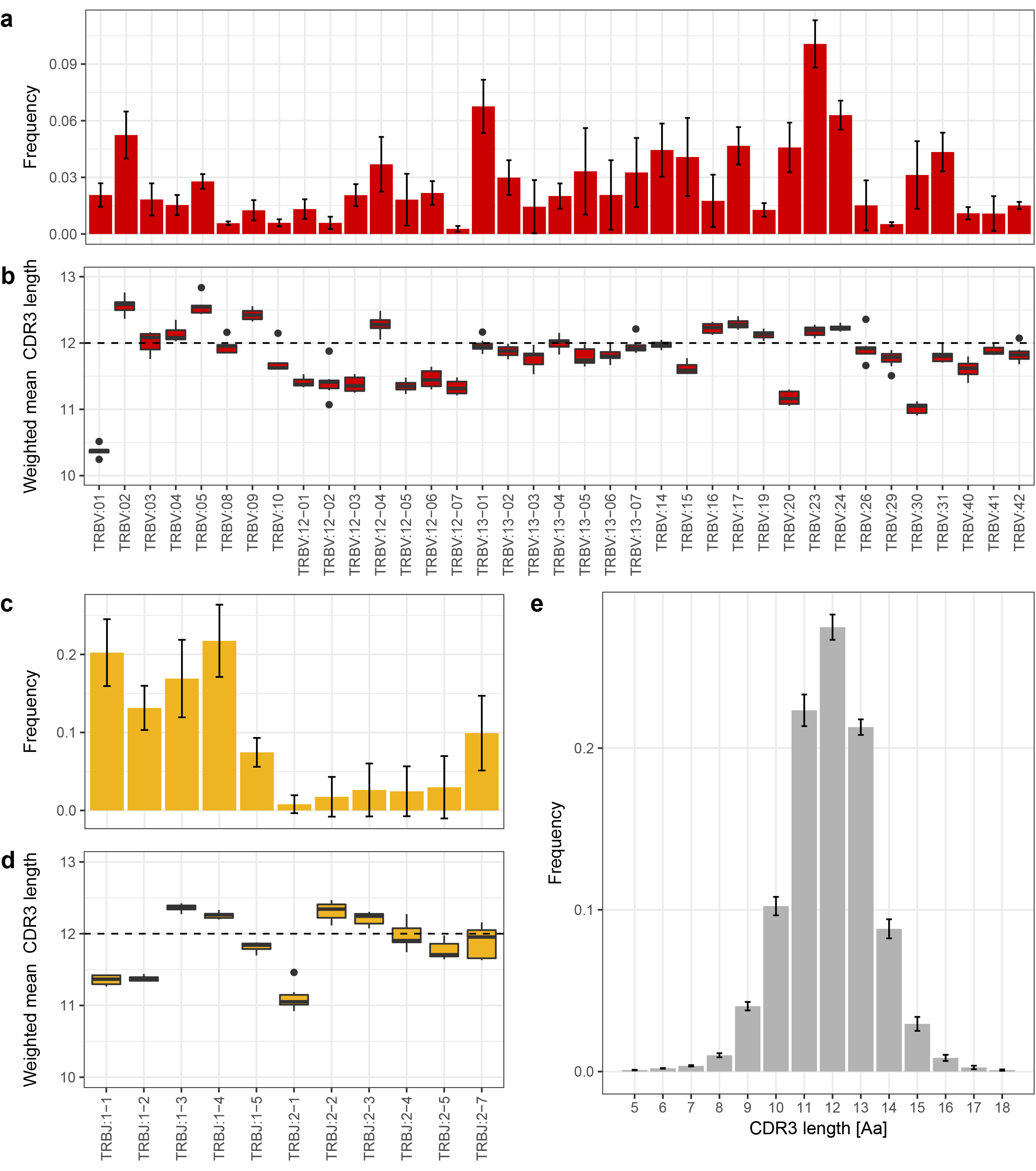
Mean V–J segment usage and CDR3 length distribution of TCRβ in seven bank voles. **(a)** Relative usage frequencies of each BV_TRBV gene. **(b)** Weighted mean (by the number of reads) of the CDR3 amino acid length for each BV_TRBV gene. **(c)** Relative usage frequencies of each BV_TRBJ gene. **(d)** Weighted mean (by the number of reads) of CDR3 amino acid length for each BV_TRBJ gene. **(e)** CDR3 amino acid length distribution. In **(a, c, e)**: the bar height indicates the respective mean frequency from seven individuals, error bars show standard deviations. In **(b, d)**: Tuckey boxplots represent weighted mean CDR3 length formed by each contributing gene segment in seven individuals; whiskers indicate data points within the 1.5 interquartile range; the dashed line marks the global CDR3 length median.

**Figure 5.**
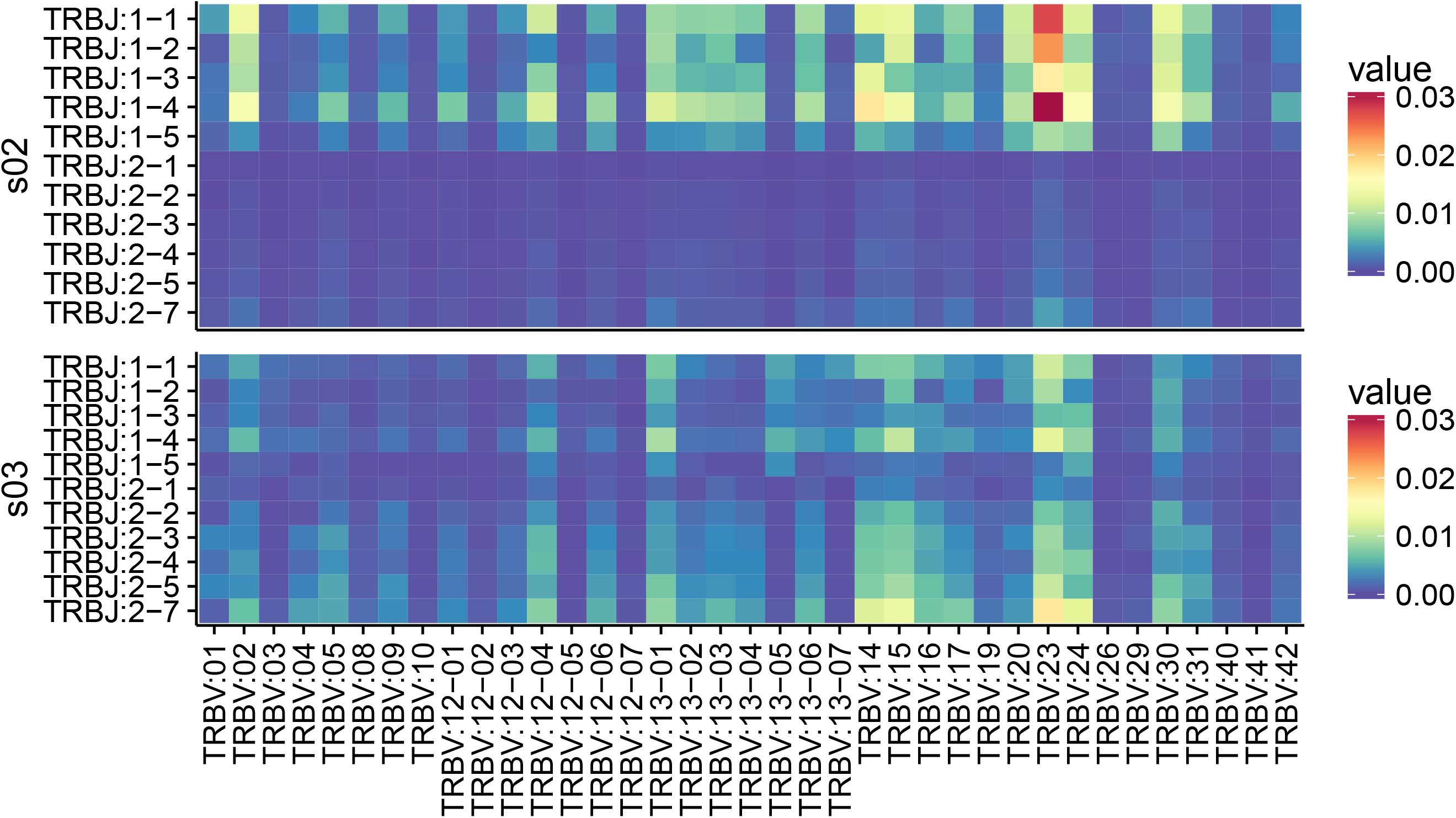
Differences in the TCRβ V–J segment paring frequencies between two individuals. Heatmap of frequencies of the CDR3 variants formed by the given V–J combinations.

The CDR3 lengths had a bell-shaped distribution and ranged from 5 to 18 amino acids (15–54 nt). The median length equals 12 amino acids, 71% of unique CDR3 sequences had a length of 11–13 amino acids, and 90% of CDR3s had length ranging from 10 to 14 amino acids (Fig. 4e). *In silico* spectratypes with V and J segments are shown in Supplementary Fig. S8. The global mean length of CDR3s was 11.91 amino acids (weighed by read count), but mean lengths of CDR3s formed by different V and J segments showed consistent differences (Figs. 4b, 4d). For example, V segments BV_TRBV01, BV_TRBV20, and BV_TRBV30 tend to form shorter CDR3s (means: 10.37, 11.17, and 11.02, respectively), whereas BV_TRBV02 and BV_TRBV05 slightly longer ones (mean 12.56, for both). Similarly, J segments BV_TRBJ1–1, BV_TRBJ1–2, and BV_TRBJ2–1 form shorter than average CDR3s (mean: 11.35, 11.37, and 11.11, respectively).

### Quantitative analysis of the bank vole TCRβ repertoire

For three additional individuals, a modified 5′ RACE protocol integrating UMIs and Illumina paired-end 150-bp HiSeq sequencing (Supplementary Fig. S1) was used to estimate the TCRβ repertoire size. The repertoire size (i.e. number of unique CDR3 variants per individual) was estimated using the Chao2 estimator, based on UMI error-corrected, unique CDR3s from four independent replicated samples per individual.

## Repertoire size estimation

The mean number of observed, unique (non-redundant) TCRβ CDR3 sequences per amplicon subsampled to 1.5 mln reads was 1.3 ×10^5^ (SD = 0.1 ×10^5^) for nucleotide sequences and 1.1 × 10^5^ (SD = 0.1 × 10^5^) for amino acid sequences (Supplementary Table S2). The mean total number of observed, unique CDR3 sequences per individual (adding up the four replicates) was 1.8×10^5^ (range: 1.5–2×10^5^) nucleotide sequences and 1.5×10^5^ amino acid sequences (range: 1.2–1.6×10^5^).

A major obstacle in repertoire-size estimation in mammals is the inability to capture the entire repertoire, even if the library from a given sample can be exhaustively sequenced^43,44^. This caveat–inherent to repertoire sampling and revealed with exhaustive sequencing experiments–results from the fact that the diversity found in any aliquot of cells will typically reflect a fraction of the total repertoire^13,43^. For that reason, richness estimators–traditionally used in ecology to tackle “unseen species” problem–are applied to calculate total clonotype diversity. They allow to estimate, from given samples, how many unobserved species (or in this case–clonotypes) are present in an individual. Following other authors^5,45^ we used a non-parametric, incidence-based estimator Chao2^46^ to estimate the lower boundary of TCRβ repertoire size. The mean of the lower-bound estimate was 2.1 ×10^5^ (range 1.7–2.3 ×10^5^) different nucleotide CDR3 sequences (Supplementary Table S3), which indicates that in four samples we directly observed up to 86% splenic TCRβ CDR3 sequences.

In our size-estimation protocol, we sub-sampled amplicons to control for differences in the sequencing depth. The 1.5 mln raw reads sufficed to retrieve most of the diversity present in each amplicon and in the individual (across all replicated amplicons). For example, for the highest-coverage individual (s08, mean: 3.5 mln raw reads per replicate), with subsampling to 1.5 mln, the unique number of observed CDR3s was 1.4×10^5^ nucleotide sequences per replicate and, altogether, 1.9× 10^5^ for the individual (when a total of 6 mln reads were analysed). Doubling the sequencing effort increased the number of unique CDR3s to 1.7×10^5^ per replicate, and to a total of 2.1 ×10^5^ for the individual (when all 13.9 mln reads were analysed). The Chao2 estimator was robust to those variations, as the estimates of CDR3 repertoire size increased only by 8%–from 2.3 ×10^5^ (at 1.5 mln reads per replicate) to 2.5 ×10^5^ (with ± 3.5 mln reads per replicate).

## Private versus public repertoire

The vast majority of CDR3 nucleotide sequences represented private TCRβ repertoire–that is, these sequences were not present in any other individual. On average, only 3.7% of CDR3 nucleotide sequences was shared between any two individuals, and 1.3% was present in all three individuals, thus representing a public repertoire (Supplementary Fig. S9). In contrast, comparison of common CDR3 nucleotide sequence between replicates from the same individual showed that, on average, 77% of sequences was shared between any two replicates, and 47% was present in all four replicates. The number of shared amino acid CDR3 sequences between any two individuals was, on average, 14.3% (SD: 1%). Further, 7.5% CDR3 amino acid sequence was present in all the individuals (Supplementary Fig. S9; Supplementary Table S3).

## Discussion

Taking advantage of advances in HTS, we were able to characterize, for the first time, the TCR repertoire of a non-model rodent. We described V and J gene segments in the bank vole, analysed their phylogenetic relationship to model murine TCRs, and also characterized basic features of its TCRβ repertoire, such as CDR3 length distribution, V–J gene usage, and a conservative estimation of the TCRβ repertoire size.

Overall, the orthology of bank vole V and J segments to mouse genes was conserved, as was the number of V and J segments. The conservation of TCR structure, manifested in the long branches of the phylogenetic trees, may result from an obligatory interaction of TCRs with MHC during antigen recognition. Olivieri *et al*. (2013) drew a similar conclusion from a comparison of V genes automatically extracted from WGAs of the mouse and rat. Their analysis showed that the phylogeny of V segments of TCRs had been characterized by much longer branches, compared to V segments of structurally similar–yet neither MHC nor otherwise restricted–immunoglobulin receptors. A constrained binding interface of co-evolving germline-encoded TCR loops and conserved features shared by all (otherwise polymorphic) MHC alleles should preserve orthologous relationship between genomic TCR segments over large evolutionary time scales^47,48^.

Despite overall conservation, a few bank vole V loci did not have a clear murine orthologue. One reason for the unclear orthology could be deletion of some genes in one of the species. For example, the bank vole BV_TRBV41 gene misses the mouse orthologue, but it closely groups with the TRBV10 gene, which is pseudogenised in mice. This suggests that genes belonging to this clade might have become non-functional in mice, and some may have been eventually deleted. Another reason could be expansion after the diversification of species. For example, TRBV12 and TRBV13 groups have multiple genes both in voles and mice. These duplicated genes group by species, which suggests that they have expanded independently, after the split. A database collecting numerous V genes extracted from WGAs (Vgenerepertoire.org^41^) shows an expansion of TCRβ V segments in many species from the Cricetidae family, with more than 30 genes found in prairie vole (*Microtus ochrogaster*) and North American deer mouse (*Peromyscus maniculatus*). But, we note that the number of genes recovered from a WGA may strongly depend on the quality of the assembly, and should be treated with caution. The WGA-based method may substantially underestimate the true diversity and expansion of V genes. In case of the bank vole, more than a half of the TCRβ V genes were missing from the automatically extracted V genes, compared to our direct sequencing of TCRβ transcripts.

Bell-shaped CDR3 length distribution, preferential use of particular V and J segments, and unequal frequencies of V–J pairings that we found are all well-documented features of the TCR repertoires of other vertebrates^10,23,49,50^. Between-individual similarities in V–J segment usage and pairing could be explained by chromatin conformation, physical proximity of germline segments, and/or recombinatorial bias^51–53^. In contrast, thymic selection, which depends upon individual MHC composition, may be responsible for the observed between-individual differences^54–56^. Another possible explanation involves clonal T cell expansion during immunization with different antigens^50^. However, standardized and controlled laboratory conditions under which the voles were kept render differential antigen exposure unlikely, thus individual skewing of their TCR repertoires most probably stemmed from some other mechanism.

A lower bound of the TCRβ repertoire size in the bank vole was conservatively estimated at 2.1 × 10^5^ unique CDR3 types, which is comparable to the estimate for the mouse, 5–8 ×10^5^ (based on Sanger sequencing of a single CDR3 length band and interpolation over the sizes of all bands^6^). Less than 4% of nucleotide CDR3 sequences was shared between any two individuals, and only 1.3% was present in all three individuals, which is similar to the numbers reported for human^43^. The number of shared amino acid sequences was almost four times greater, and is of the same order of magnitude as found for human and mice^43,57^. We note that, although the number of observed sequences was not far from the estimated repertoire (on average, 86%), we do not have full individual repertoires and the number of shared clonotypes may be underestimated. However, it was previously shown that the frequency of a given TCR type (within-sample abundance or presence in multiple aliquots from a single individual) predicts sharing of this clonotype between individuals^58^. Therefore, while more exhaustive repertoire sequencing should identify less frequent variants, it would be unlikely to significantly increase the size of the shared and public repertoire.

Taken together, our work broadens knowledge of the immune system of the bank vole-a popular species in ecological and evolutionary research. We showed that an informative analysis of an individual TCR repertoire is possible with partial information on the genomic composition of the TCR locus (e.g., we did not have full sequences of genomic segments prior to somatic recombination and had only provisional assignment to the actual loci for V and J segments). In the future, the methodology developed and tested herein can be used to study immune response in nature and in experiments involving this species. More importantly, our TCR profiling workflow can be easily adapted to other non-model vertebrates that lack genetic resources (e.g. well-annotated reference genomes or V/J segment listings in bioinformatics databases), do not have dedicated molecular reagents and kits (i.e. monoclonal antibodies), and cannot be analysed with human or mouse-specific software, which relies on published references of genomic segment. The 5′ RACE-based protocol requires only one species-specific set of primers in the constant region of the TCR molecule, which is easy to obtain. Effective primer design is possible based on the *de novo* draft genome or transcriptome assemblies of this region, which are relatively easy and straightforward as compared to assemblies of highly duplicated and diversified V and J segments. The data obtained can be analysed with our publicly available AmpliTCR and AmpliCDR3 programs, which extract V and J segments and the CDR3 region based on conserved amino acid residues. Therefore, with a rapid increase in the number of genomes and transcriptomes available, the promise of *“no more non model species*” in the HTS era^59^ can be finally fulfilled for the challenging task of immune repertoire sequencing. We hope that our approach will become particularly useful in ecological and evolutionary research, where organisms of interest rarely belong to the exclusive club of well-established model species. Yet, there is great potential in such studies–from enhanced comparative immunology to capability of tracking immune response and disease progression in endangered vertebrate species.

## Materials and methods

### Primer design

The bank vole is a non-model organism, with limited genetic information available. To design nested primers for 5′ RACE, we used *de novo* assembled transcriptomes from spleens of seven animals. For a detailed description of the dataset and the assembly strategy, see Migalska *et al*.^37^ Briefly, transcriptomes were *de novo* assembled with Trinity^60^, and contigs highly similar to the first exon of the constant regions of TCRβ *Mus musculus* sequences (as extracted from Genbank records: AH002088.2, AH002089.2, and X03574.1) were retrieved. Based on these contigs, three nested primers in the 3′ end of the constant region were designed: MyglTCRb_1: TGATCTCTGCTTCTGATG, MyglTCRb_2: GATGGCTCAAACAGGGTGACC, and MyglTCRb_3: GGACTCACCTTGCTCAGATCCT. The full list of primers and adaptors used in the study is given in the Supplementary Table S4, and a schematic illustration of the primers and adaptor location is presented in the Supplementary Fig. S1.

### Experimental animals and RNA extraction

Spleens from healthy laboratory bank voles (n = 10) were collected during necropsy in accordance with internationally recognized guidelines for research on animals approved by the Krakow Ethical Committee for Experiments on Animals. Information on the animals’ sex, age at death, as well as on its assignment to the qualitative or quantitative part of the study is given in Supplementary Table S5. Fragmented spleens (4–6 fragments depending on spleen size) were preserved in RNAlater (Sigma-Aldrich), and homogenized piece by piece with FastPrep® (MP). Total RNA was extracted using RNAzol® RT (Sigma-Aldrich) according to the manufacturer’s instructions, and eluted in 50 μl RNase-free water. For qualitative description, seven bank voles were used (one spleen fragment per individual). For quantitative analysis of three additional bank voles, aliquots (20 μl) of extracts from each spleen fragment were pooled and purified on NEXTflex™ Poly(A) Beads (Bioo Scientific®) to remove abundant rRNA and reduce sample volume (elution in 15 μl).

### 5′ RACE-based library construction and Illumina sequencing

For **qualitative description** of the bank vole TCRβ repertoire, we used the 5′ RACE method, based on the procedure described by Mamedov and colleagues (2013), but murine-specific primers were replaced with bank vole-specific ones (see *Primer design* section). cDNA synthesis was conducted using the 5′ template-switching procedure, followed by nested PCR amplification steps, described in detail in Mamedov *et al*.^14^, and summarized in the Supplementary Fig. S1. For the first step (cDNA synthesis), the Mint-2 cDNA synthesis kit (Evrogen) containing the 5′ oligo adaptor PlugOligo-1 was used, with custom-made MyglTCRb_1 primer (1 μM) and 2 μg total RNA for each sample. Subsequent PCRs were conducted using the high-fidelity Encyclo polymerase mix (Evrogen) according to the manufacturer’s instructions. In the first PCR, 1 μl of the first strand cDNA was used as a template, with Smart20 oligonucleotide and nested MyglTCRb_2 primer (Supplementary Table S4). In the second PCR, 1 μl of the product of the first-round reaction was used a as a template, with Step_1 oligonucleotide and MyglTCRb_3 nested primer (Supplementary Table S4). PCR conditions for the first PCR were as follow: 1-min denaturation at 95°C, 20 cycles of 95°C for 20 s, 65°C for 20 s, and 72°C for 50 s, with a final elongation for 3 min at 72°C. Conditions of second PCR were the same, except for the cycles (their number was reduced to 10) and final elongation prolonged to 5 min. PCR products were run on 1.5% agarose gel, and bands of desired size (~600 bp) were excised using QIAquick Gel Extraction Kit (QIAGEN) according to manufacturer’s protocol. The final library was constructed with NEBNext® DNA Library Prep Master Mix Set for Illumina® (NEB). Paired-end (PR), 300-bp sequencing was performed at the Institute of Environmental Sciences, Jagiellonian University, Krakow (Poland), with the MiSeq Reagent Kit v3 on a MiSeq sequencer (Illumina). The sequencing yielded more than 20 mln raw reads (mean per sample: 2.9 mln, SD: 0.2 mln; Supplementary Table S6).

The protocol was modified for **quantitative analysis** to incorporate UMIs during cDNA synthesis^18^, such that a cDNA strand synthetized from a single mRNA molecule is uniquely marked. These UMIs (tags composed of 12 random nucleotides) enable more confident discrimination of sequencing errors from biological variants. During the cDNA synthesis, PlugOligo-1 was replaced with SmartNNNNa 5′ adaptor which contained dU nucleotides and a UMI^18^. Reactions were conducted with Mint-2 cDNA synthesis kit (Evrogen), with a MyglTCRb_1 primer, and ~30 ng purified mRNA for each sample. cDNA synthesis was conducted at 42°C for 45 min, and for an additional 1.5 h after the addition of IP solution (5μl/sample) for an additional 1.5h. Products of cDNA synthesis were treated with 5U USER enzyme (uracil–DNA glycosylase, NEB) for 1 h to degrade any leftover SmartNNNNa adaptor. Moreover, amplification steps were modified to avoid sample-consuming adaptor ligation in the final phase of the library preparation. Instead, a two-step PCR-based primer extension method was used, with modified primers containing partial Illumina adaptor sequences (Supplementary Table S4). This method was proven to be superior to adaptor ligation in terms of yield and efficiency in the immune repertoire library preparation^61^. For each individual, there was a single first-step PCR and four, independently tagged second-step PCRs. In the first PCR (50 μl volume), 5 μl first-strand cDNA was used as a template with 0.5 μM of primers Smart20-mod and TCRb_3NN (Supplementary Table S4). Subsequently, samples were purified using Agencourt AMPure XP beads with 1:0.6 ratio of DNA:beads, to remove short, nonspecific products and primer/adaptor-dimers. Then, 20 μl of purified eluate was separated into four parts for second PCRs (25 μl volume each) with 1.5 μM of specific P5_50X/P7_70X primer combinations (Supplementary Table S4), thus allowing for de-multiplexing of samples pooled in a sequencing run. PCR conditions were as described previously, with a modification of number of cycles (first PCR: 23 cycles, second PCR: 14 cycles). Products of second PCR were run on 1.5% agarose gel, and bands of desired size (~600 bp) were excised using the QIAquick Gel Extraction Kit (QIAGEN) according to manufacturer‘s protocol. The final library containing 12 amplicons (three individuals in four replicates) was sequenced by Macrogene (Seul, Korea) as a part of the 150-bp PE sequencing run on the Illumina HiSeq 2500 instrument. The sequencing yielded 1.5–4.3 mln raw PE reads per amplicon (mean: 2.8 mln per sample, Supplementary Table S2).

### Qualitative description of the bank vole TCRβ repertoire

The TCRβ repertoire for seven individuals (s01–s07) was analysed and described, as detailed in the following steps: (i) pre-processing of paired-end Illumina reads; (ii) read sub-sampling, extraction of V and J segments and retrieval of consensus V and J variants by AmpliTCR; (iii) phylogenetic analysis and naming of the V and J variants; and (iv) description of the V–J segment usage and CDR3 length distribution with AmpliCDR3.

All bioinformatics tools used in the following sections (AmpliMERGE, AmpliCLEAN, AmpliTCR, and AmpliCDR3) belong to the AmpliSAS^62^ family within the AmpliSAT suite (Amplicon Sequencing Analysis Tools) available at: http://evobiolab.biol.amu.edu.pl/amplisat/.

## Pre-processing of Illumina data

Overlapping paired-end reads from the 5′ RACE-based library (MiSeq 2×300 bp run) were merged with AmpliMERGE–a tool based on FLASH^63^ –using default parameters. Subset of 2 million randomly chosen, merged reads was filtered with AmpliCLEAN, removing reads with lower average Phred quality score (<30) and those not matching the last 12 nucleotides of the MyglTCRb_3 primer (Supplementary Table S4) at the beginning of the constant TCRβ region. On average, 1.8 mln reads per sample passed the pre-processing step (Supplementary Table S6).

## Read sub-sampling, TCRβ V and J segments extraction, and retrieval of consensus V and J variants by AmpliTCR

To analyse a comparable number of TCRβ sequences for each individual, 1 mln random reads per sample were selected for further processing; 95% of these reads had lengths between 325 and 536 bps and covered the full variable region of the TCRβ, including the rearranged and spliced V(D)J segments. Identification of V and J segments was performed automatically by AmpliTCR, whereas the D segment was neither extracted nor identified, as too much uncertainty about its germline sequences is introduced by N-diversity regions. The AmpliTCR uses protein patterns in the PROSITE format^64^ to identify highly conserved residue positions in the translated sequence. To extract TCRβ V segment (TRBV) the following pattern was used: x(5)-Q-x-P-x(14)-C-x(10,11)-W-Y-x(39,42)-[LM]-x(14)-C-x; and for TCRβ J segment (TRBJ): x(4)-G-x-G-x(2)-L-x-[VI]-x. Patterns were composed of conserved amino acids previously described in the literature^65,66^ as well as the multiple alignment of human and mouse TRBV and TRBJ sequences found in the IMGT database^67^. On average, 91% of the analysed reads (range: 87%–94%) contained both protein patterns. AmpliTCR discards sequences containing unspliced introns, frameshift indels, and premature termination codons (PTCs). Approximately 41% of TCRβ sequences with recognized protein patterns were later discarded for those reasons.

Extracted TRBV and TRBJ sequences were de-replicated by AmpliTCR into unique, non-redundant variants and ordered by sequencing depths. Next, TRBV variants with frequency less than 0.01% were removed to discard artefacts, such as sequencing errors, PCR-based substitutions, and chimeras. For TRBJ, the frequency threshold was increased to 0.1%. Both frequency values are based on the minimum percentage of rare alleles for each TCRβ segment found in human^68^. Further, to remove the remaining artefacts, variants were clustered with the CD-HIT-EST^69^ algorithm (implemented in AmpliTCR), with a minimum identity threshold of 95% for TRBV and 85% for TRBJ. These thresholds were established based on sequence similarity between the TRBV and TRBJ loci in human and mouse^70^. In addition, to prevent merging true but similar variants, sequences with within-cluster frequencies above 20% were moved to form a new cluster. Finally, major variants within each cluster were selected as putative TRBV and TRBJ alleles. All alleles are available in the Supplementary Files S1 and S2.

## Phylogenetic analysis and naming of the V and J segments

No genomic information about segments composing bank vole TCRβ was available prior to the analysis; therefore, provisional loci identification was conducted based on both orthology (co-occurrence of bank vole alleles and murine reference within the same phylogenetic clade) and sequence similarity to the mouse references from the IMGT database^2^ (the full list of the murine reference sequences with accession numbers is available in a Supplementary Table S1, all retrieved bank vole alleles are in the Supplementary Files S1 and S2). The orthology was assessed from phylogenetic trees constructed separately for all V and J alleles in MEGA6^71^. The neighbour-joining trees were constructed with the evolutionary distances computed using the Maximum Composite Likelihood method, with a bootstrap of 1000 replicates (Supplementary figures S2 and S3). In case of J segments, several branches were poorly resolved, with low bootstrap values. This is most likely caused by the short length of sequences (<25 bp), which resulted from trimming them at the border of the CDR3 region. However, visual inspection of an alignment of the bank vole sequences to the murine reference allowed confident loci assignment. We named bank vole loci (clades of putative alleles) with a “BV” prefix (BV_TRBV for V segments and BV_TRBJ for J segments), while keeping the numbering of the murine orthologues. Alleles within each locus were numbered arbitrarily. A few V segment sequences did not have a close murine ortholog (or a highly similar sequence); therefore, to avoid confusion, their clusters were named with BV_TRBV**4**X (as none of the murine V segments has a number of 40 or higher). Supplementary phylogenetic analysis was conducted for V segments, with the addition of genes automatically extracted from the bank vole draft genome (GenBank assembly ID: LIPI00000000.1) and stored in an online database Vgenerepertoire.org^41^ (accessed: May 20, 2017).

## V–J segment usage and CDR3 length distribution

After the V and J segment loci were identified and named, we assessed V–J segment usage and CDR3 length distribution in the seven bank vole individuals. Pre-processed Illumina reads (see above) were re-analysed with AmpliCDR3 tool, with the reference BV_TRBV and BV_TRBJ segments obtained in the previous steps. For each individual V and J segment, sequences are locally aligned with BOWTIE2^72^ (--sensitive-local -k 2) to the BV_TRBV/J references. Particular BV_TRBV and BV_TRBJ loci were annotated if they had aligned with more than 90% identity to any of the reference sequences provided.

The CDR3 region is defined as the sequence between the last conserved Cys in the V segment and one residue (3 nt) before the first conserved Gly in the J segment. AmpliCDR3extracts the CDR3 region by searching for a TG[TC][GA][CG] motif located at the 3’ end of the V segment and a ([GA][CG]) motif marking the 5′ beginning of the CDR3 region. The 5′ end of the CDR3 is fixed at 31-bp upstream from the 5′ end of the constant TCRβ region (sequence of the MyglTCRb_3 primer, Supplementary Table 4, Supplementary Fig. 1). AmpliCDR3 identified the correct protein pattern for CDR3 regions in approximately 55% of the pre-processed reads (44%–63%). Finally, the AmpliCDR3 filters putatively non-functional CDR3 sequences (shorter than 15 bp and longer than 54 bp, containing frameshift indels and PTCs). After filtering, on average, 85% of the reads were retained (75%-89%). Although the remaining sequences likely still contain substitution errors introduced during PCR or sequencing, they should not significantly affect the descriptive characterization of the TCRβ repertoire (i.e. V–J segment usage and CDR3 length distribution). Qualitative statistics were calculated and visualized with R^73^.

### Quantitative analysis of the bank vole TCRβ repertoire

For three additional individuals (s08–s10), a quantitative analysis of the TCRβ CDR3 repertoire was conducted with a UMI-based error-correction strategy. Data were analysed with the tools AmpliMERGE, AmpliCLEAN, and AmpliCDR3 in the following steps: (i) pre-processing of paired-end Illumina reads; (ii) sub-sampling, identification of CDR3 sequences and filtering; (iii) UMI-based CDR3 error correction; (iv) repertoire size estimation; and (v) private versus public repertoire comparison.

## Pre-processing of Illumina data

Non-overlapping, paired-end reads from Illumina HiSeq2500 (2×150 bp) were concatenated with AmpliMERGE and filtered with AmpliCLEAN to remove reads with average Phred quality score lower than 30. The reads did not cover the whole variable region of the TCRβ chain (Supplementary Fig. S1), but retrieved the UMI, the entire CDR3 and J segment, as well as the partial V segment.

## Read sub-sampling, CDR3 region identification, and filtering with AmpliCDR3

To control for differences in sequencing depth that might influence the number of retrieved variants, we subsampled pre-processed reads up to 1.5 mln. Rarefaction curves analysis showed that, at this point, the number of new CDR3 sequences found within an amplicon started to plateau, such that our sub-samples likely captured most of the CDR3 diversity within an amplicon (Supplementary Fig. S10).

CDR3 region identification and filtering was undertaken with AmpliCDR3, as described in the subsection on *V-J segment usage and CDR3 length distribution*. Conserved protein motives delimiting CDR3 were found in 95% (SD = 3%) of the reads; on average, 72% of them (SD = 5%) passed the filtering criteria (Supplementary Table S2).

## UMI-based CDR3 error correction

For reliable estimation of the number of unique TCRβ CDR3 variants within an amplicon, we conducted UMI-based error correction. As UMIs are added to molecules during cDNA synthesis, all HTS reads sharing identical UMIs should originate from the same, amplified cDNA template. Any substitution or indel among reads with identical UMI can, therefore, be identified as an artefact. At the same time, any two sequences tagged with different UMIs can be inferred to represent true, biological variants even if they differ by as little as a single nucleotide substitution^18^. Our error-correction algorithm, implemented in AmpliCDR3 tool, is based on the principles of the MIGEC (molecular identifier groups–based error correction) strategy developed by Shugay and colleagues (2014), but it is simplified to increase speed and modified to account for lower sequencing depths. To infer the correct sequence of a CDR3, AmpliCDR3 groups reads with identical UMIs into molecular identifier groups (MIGs)^18^. As the majority of errors arise during late PCR cycles or sequencing, they are expected to represent the minority of reads within a MIG, and a consensus sequence is inferred to be the true CDR3 sequence. However, early-stage PCR errors, presence of PCR chimeras, and so on can produce ambiguous, “mosaic” MIGs. For that reason, the AmpliCDR3 inspects each MIG and discards those in which half of the reads fails to pass the similarity threshold (2-bp substitution) to the most abundant sequence (thereby 0.6% of reads were discarded). If a MIG contained only two reads, they had to be identical to remain in the analysis as a true sequence. MIGs that contained only one read were discarded (29%). Finally, identical CDR3s with different UMIs might derive from one T cell (multiple mRNA transcripts), or from different T cells bearing an identical CDR3 part of the TCR. Regardless of the source, only unique (non-redundant) CDR3 variants were counted. Lastly, CDR3 nucleotide sequences were translated into amino acid sequences.

## Repertoire size estimation

To estimate the lower bound of individual TCRβ diversity (repertoire size) we chose a non-parametric, incidence-based Chao2 estimator^46^. Chao2 does not require assumption of a priori clonotype distributions, and it had been previously applied to TCR repertoire estimation^5,45^. In contrast to abundance-based estimators, which use absolute counts of T cells bearing the given CDR3 (clonotype), Chao2 uses presence–absence data across replicates. We preferred to avoid abundance-based estimators as it is difficult to infer absolute T cell numbers (i.e. abundance of a given clonotype) by counting the observed mRNA transcripts. The reason is that a differentiation between identical CDR3 sequences extracted from mRNAs coming from a single T cell and those from different cells but bearing identical CDR3s (i.e. expanded clones or instances of convergent recombination) is impossible. For calculations, we used the following equation (1):

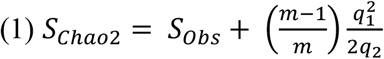

where *S_obs_* is the total number of CDR3 unique sequences observed across all replicates for a given individual, *q_1_* is the number of CDR3 sequences present only in one replicate, *q_2_* is the number of CDR3 sequences present in two replicates, and *m* is the number of replicates (m=4). Calculation of CDR3 sequences [nt] present in one, two, three or all four replicates for each individual was conducted automatically by AmpliCDR3.

## Private versus public repertoire

To compare TCR diversity among individuals, we first pooled unique CDR3 sequences from four replicates for each individual and removed redundant sequences. The resulting repertoire was compared among individuals–both at the nucleotide and amino acid levels. We further defined a CDR3 sequence as “private” if it was present in only one bank vole in our analysis, and as “public” if it was shared by all three individuals. Calculations were conducted with AmpliCDR3.

## Data accessibility

Illumina MiSeq and HiSeq raw reads (see the *Materials and Methods* section) are deposited in FASTQ format in the European Nucleotide Archive (ENA), with the study accession PRJEB22487. V and J segment consensus alleles are available in the Supplementary Materials associated with this article. Bioinformatics tools AmpliTCR, AmpiCDR3, AmpliMERGE, and AmpliCLEAN are available for download or online use at our laboratory server: http://evobiolab.biol.amu.edu.pl/amplisat/.

## Acknowledgments

The authors sincerely thank P. Koteja for the donation of bank vole spleens. The authors also thank W. Babik and K. Dudek for running the Illumina MiSeq sequencing and M. Koczal for comments of an earlier version of the manuscript. This work was supported by the National Science Centre (NCN) grant UMO-2013/08/A/NZ8/00153 to J. Radwan; M. Migalska received a PhD scholarship UMO-2017/24/T/NZ8/00088 from the NCN.

## Author Contributions

M.M. and J.R. designed the experiments, M.M. conducted the experiments, A.S. wrote the AmpliSAT suite, A.S. and M.M. conducted the bioinformatics analysis, and M.M., A.S, and J.R. analysed results and wrote the manuscript.

